# Novel microRNAs downregulated in breast cancer tumors bind to the 3’UTR of SNAIL, SLUG, ZEB1 and/or TWIST and decrease metastatic behavior in breast cancer cells

**DOI:** 10.1101/2023.02.03.526978

**Authors:** Elisa Pérez-Moreno, Victoria Ortega-Hernández, Valentina A Zavala, Jorge Gamboa, Wanda Fernández, Pilar Carvallo

## Abstract

Metastasis, the leading cause of cancer-associated deaths, is promoted by transcription factors SNAIL, SLUG, ZEB1 and TWIST through the activation of epithelial-mesenchymal transition (EMT). MicroRNAs can suppress EMT, emerging as candidate molecular biomarkers and novel therapeutic targets. Herein, we evaluated microRNAs downregulated in breast cancer tissues expressing EMT transcription factors, to find new potential regulators of EMT. MiR-30a, miR-1271, miR-196a, miR-202, miR-210, miR-22, miR-331 and miR-34b were validated. Seven microRNAs downregulated luciferase activity through EMT transcription factors 3’UTR, and all microRNAs decreased cell migration, invasion and/or proliferation. In MDA-MB-231 cells, miR-196a and miR-22 decreased endogenous ZEB1 levels, and miR-30a endogenous CCR7 levels. These results suggest that microRNAs studied are novel regulators of EMT through the control of SNAIL, SLUG, ZEB1 and TWIST. They also regulate the metastatic behavior of cancer cells, and may control the development of lymph node metastasis through the regulation of CCR7.

**Graphical abstract:** 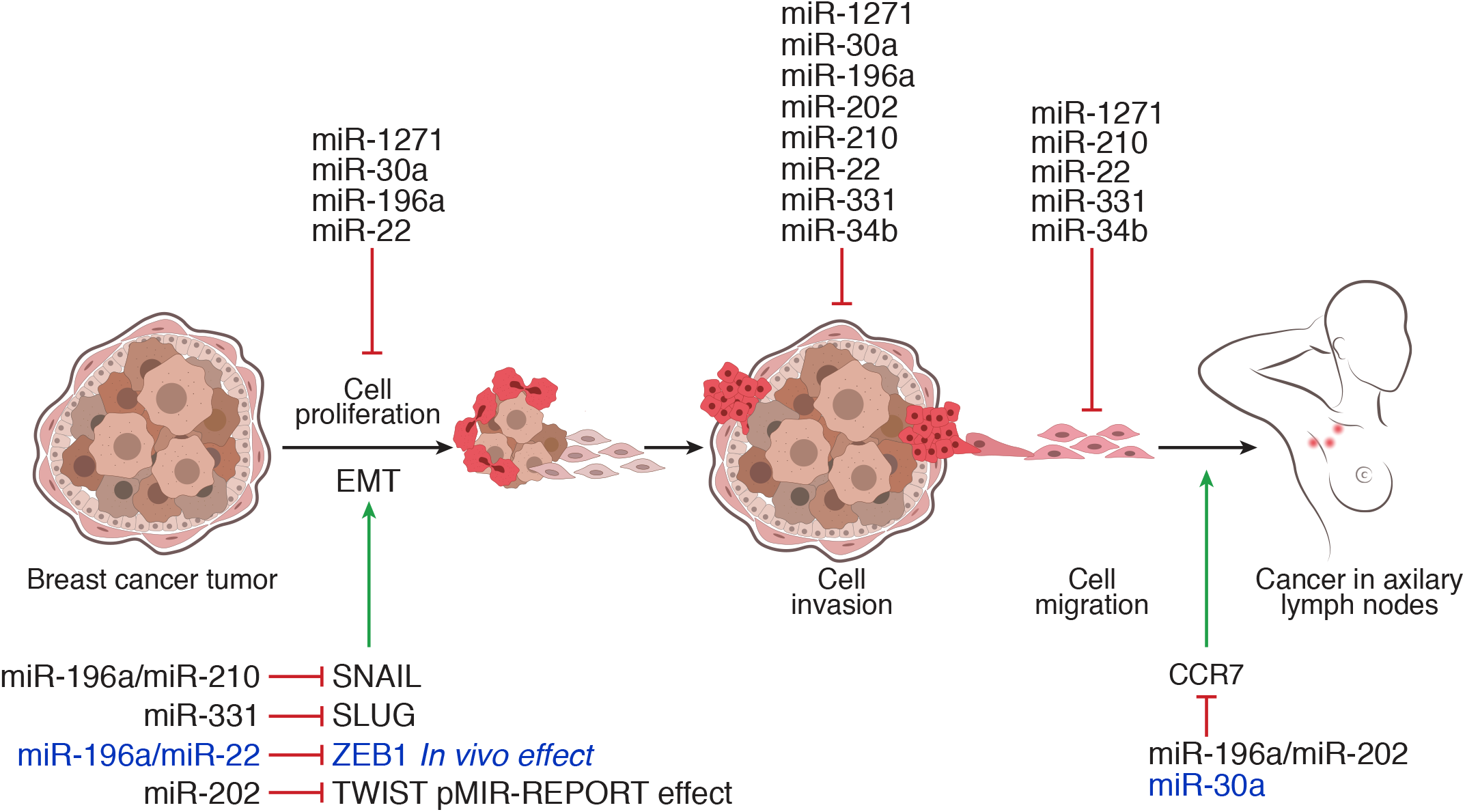

## 1. Introduction

Breast cancer is one of the main causes of death in women when it goes through a process of invasion and metastasis. Lymph nodes close to the primary breast cancer tumor have a high chance to be colonized by cancer cells leading to metastatic spots and a poor prognosis for the patient [1-3]. Therefore, metastatic process has been widely studied for years and its inhibition has been a major objective in breast cancer treatment. Epithelial-mesenchymal transition (EMT) has been studied as part of the metastatic process, in several cancer cell lines and animal models. It has been shown that EMT is mainly promoted by the transcription factors SNAIL, SLUG, ZEB1 and TWIST [4-6], enhancing the metastatic ability of cancer cells [7-11]. In human cancer tumors, there are strong evidence that an EMT-like process is occurring since SNAIL, SLUG, ZEB1 and TWIST expression has been associated with the presence of lymph node metastasis in patients [12-15]. In addition, diverse studies have associated the chemokine receptor CCR7 to lymph node metastasis, finding that its expression induces targeted migration of breast cancer cells to lymph nodes and promotes EMT in cancer cell lines [16].

MicroRNAs are small non-coding RNAs involved in metastasis development due to their role as metastatic promoters or inhibitors through the regulation of epithelial-mesenchymal transition [17]. One of the most studied is miR-200 family, playing a fundamental role in the maintenance of the epithelial phenotype by regulating EMT relevant actors like SNAIL and ZEB1 [18-19]. Together with miR-200, other microRNAs such as miR-10b, let-7 and miR-21 have a relevant participation in metastasis development [20-22]. Although there have been advances in identifying targets of microRNAs that participate in distant metastasis [23], to our knowledge, no targets have been identified for microRNAs involved in lymph node metastasis [24-27]. From the clinical point of view, the identification of microRNAs regulating specific targets, as EMT transcription factors, and therefore modulating some steps of metastasis, is highly relevant to design future therapies to improve prognosis in cancer patients.

In this study, we describe microRNAs able to regulate luciferase activity through the 3’UTR of SNAIL, SLUG, ZEB1 and TWIST. In addition, all selected microRNAs decreased migration, invasion, and cell proliferation in a metastatic cell model. MicroRNAs miR-196a and miR-22 decreased endogenous ZEB1 levels in MDA-MB-231 cells, confirming their role as new ZEB1 regulators.

## 2. Methods

### 2.1. Tissue samples

A total of 101 FFPE unselected breast cancer tumors were collected from patients prior to chemotherapy or radiotherapy, from two hospital centers of Santiago of Chile. All patients signed an informed consent, and this protocol was approved by the Ethics Committee of the “Servicio de Salud Metroplitano Central”. A pathologist previously determined histological type, tumor grade and hormonal receptor status. The clinicopathological features of the tumor samples are listed in Supplementary Table 1.

### 2.2. Immunohistochemistry in breast cancer tissues

Immunohistochemistry was performed in tumors disposed in tissue microarrays as previously described [28]. The following primary antibodies were used: SNAIL (Cell Signaling #3879), SLUG (Cell Signaling #9585), ZEB1 (Cell Signaling #3396), TWIST (Abcam ab50887), and CCR7 (Abcam ab32527). For each tumor, the H-score was calculated.

### 2.3. Selection of microRNAs for experimental validation

Candidate microRNAs were selected from microRNA microarrays previously performed in 50 breast tumors from our cohort (Zavala et al, manuscript in preparation, GEO GSE144405). RNA22 and TargetScan were used for *in silico* analysis to identify potential microRNAs as regulators of the EMT transcription factors. To select microRNAs for experimental validation we considered three parameters: 1) downregulation in tumors with expression of the EMT transcription factors, 2) downregulation compared to normal breast, 3) the potential ability to regulate more than one transcription factor. Six microRNAs were selected: miR-196a-5p for SNAIL and ZEB1, miR-202-3p and miR-34b for TWIST, miR-210-3p for SNAIL and SLUG, miR-22-3p for ZEB1 and miR-331-3p for SLUG. In addition, two previously validated microRNAs were used as positive controls: miR-30a-5p for SNAIL and SLUG, and miR-1271-5p for ZEB1 and TWIST [29-32].

### 2.4. Cell culture and transfections

Metastatic breast cancer cell line MDA-MB-231 and HEK-293T cells were maintained in DMEM/F12 medium (Gibco, ThermoFisher Scientific), supplemented with 10% FSB and 1% penicillin/streptomycin, at 37°C and 5% CO_2_. Transient transfections were carried out with mirVana microRNA mimics (Ambion, ThermoFisher Scientific), using Lipofectamine RNAiMAX Transfection Reagent (Invitrogen, ThermoFisher Scientific), according to the manufacturer’s instructions.

### 2.5. Luciferase reporter assay

The 3’UTR region of human SNAIL, SLUG, ZEB1, TWIST and CCR7 mRNAs were amplified by PCR from control genomic DNA and inserted in the multiple cloning site of the pMIR-REPORT luciferase vector (Invitrogen, ThermoFisher Scientific). HEK-293T cells were co-transfected with pMIR-REPORT luciferase plasmid and microRNA mimics at a final concentration of 50nM. Luciferase activity was detected using Luciferase Assay System (Promega), and normalized to β-gal activity.

### 2.6. Cell migration and cell invasion assays

For migration assays, 3×10^4^ transfected MDA-MB-231 cells were seeded in the top chamber of non-coated membranes (24-well insert; pore size 8 µm; SPL Life Sciences) using serum-free DMEM/F12 medium. Invasion assays were performed equally, using Matrigel coated membranes (BD Bioscience). Medium supplemented with 10% FSB was used as chemoattractant in the lower chamber. After incubation, cells that did not migrate/invade through the membrane were removed using a cotton swab. Cells attached to the lower surface of the membrane were fixed with methanol, and stained with crystal violet. Photos of five random regions were captured, and cells were counted using ImageJ.

### 2.7. Colony formation assay

Two-thousand previously transfected MDA-MB-231 cells were plated and growth in 3.5 cm wells, with complete medium. After 10 days, the cells were carefully washed with PBS 1X, fixed with methanol, and stained with crystal violet. Colonies were observed under the microscope and counted.

### 2.8. Immunofluorescence in MDA-MB-231 cells

MDA-MB-231 cells were plated over glass coverslips, and microRNA transfection was performed as previously mentioned. Cells were washed with PBS 1X, and fixed with 4% paraformaldehyde. Cells were permeabilized with Triton X-100 0.1% and blocked with 10% goat serum. Incubation with Ki67 antibody (Cell signaling #9129), nuclei staining and sample mounting was done as previously described [28]. Three zones of each coverslip were photographed and considered for the analysis.

### 2.9. Western blot in MDA-MB-231 cells

MDA-MB-231 cells previously transfected with microRNAs were lysed using RIPA lysis buffer supplemented with protease inhibitors. Total cell extracts (50 µg) were separated in 6%-12% gradient polyacrylamide gels and transferred to PVDF membranes. Membranes were incubated with antibodies against SNAIL (Cell Signaling), SLUG (Cell Signaling, ZEB1 (Santa Cruz Biotechnology), TWIST (Abcam), CCR7 (Abcam), β -actin (Santa Cruz), anti-mouse (Jackson ImmunoResearch) and anti-rabbit (Santa Cruz Biotechnology). Protein detection was assessed using the SuperSignal™ West Pico PLUS Chemiluminescent Substrate (ThermoFisher Scientific).

### 2.10. Statistical analysis

Analyses were performed using GraphPad Prism 6 software. Data is presented as the mean ± standard deviation. Student’s t test was used for comparing two groups, and one-way ANOVA and Dunnett test for comparisons between different groups. Fisher’s exact test and chi-squared test were used to analyze contingency tables. Results were considered statistically significant when p<0.05.

## 3. Results

### 3.1. Expression of SNAIL, SLUG, ZEB1 and TWIST in breast cancer

The expression of the transcription factors SNAIL, SLUG, ZEB1 and TWIST has been detected in 53/101 primary breast cancer tumors. Each transcription factor was detected at least in one tumor (Supplementary Fig.1A), being ZEB1 the most frequently expressed (36/53 tumors, 68%), followed by SNAIL (21/53 tumors, 40%), SLUG (12/53 tumors, 22.6%) and TWIST (11/53 tumors, 20.8%). SLUG expression showed association with tumor grade (p=0.0039), and TWIST with tumor size (p=0.0241), in concordance to previous reports [13,15]. No association between the expression of EMT transcription factors and lymph node metastasis was found.

### 3.2. Selected microRNAs regulate luciferase translation trough the 3’UTR of EMT transcription factors

We selected six microRNAs as candidate regulators of EMT transcription factors expression. Candidate microRNAs were first validated through luciferase reporter gene assays. As expected, the two positive control microRNAs selected for this assay (miR-30a and miR-1271) reduced the luciferase activity in the presence of their reported targets when compared to a negative control microRNA (miR-NC) (Fig.1A-D). From the candidate microRNAs, miR-196a, miR-210, miR-202, miR-22 and miR-331 decreased luciferase activity through their predicted 3’UTR targets, suggesting that these microRNAs may suppress the expression *in vivo* of the EMT transcription factors. Only miR-34b failed to decrease luciferase activity through its predicted 3’UTR target, discarding it as a direct regulator of TWIST expression (Fig.1D). These results reveal that miR-196a, miR-210, miR-331, miR-22 and miR-202 interact with their predicted 3’UTR targets downregulating protein synthesis, suggesting that these microRNAs are good candidates to regulate EMT *in vivo*.

**Fig 1.**
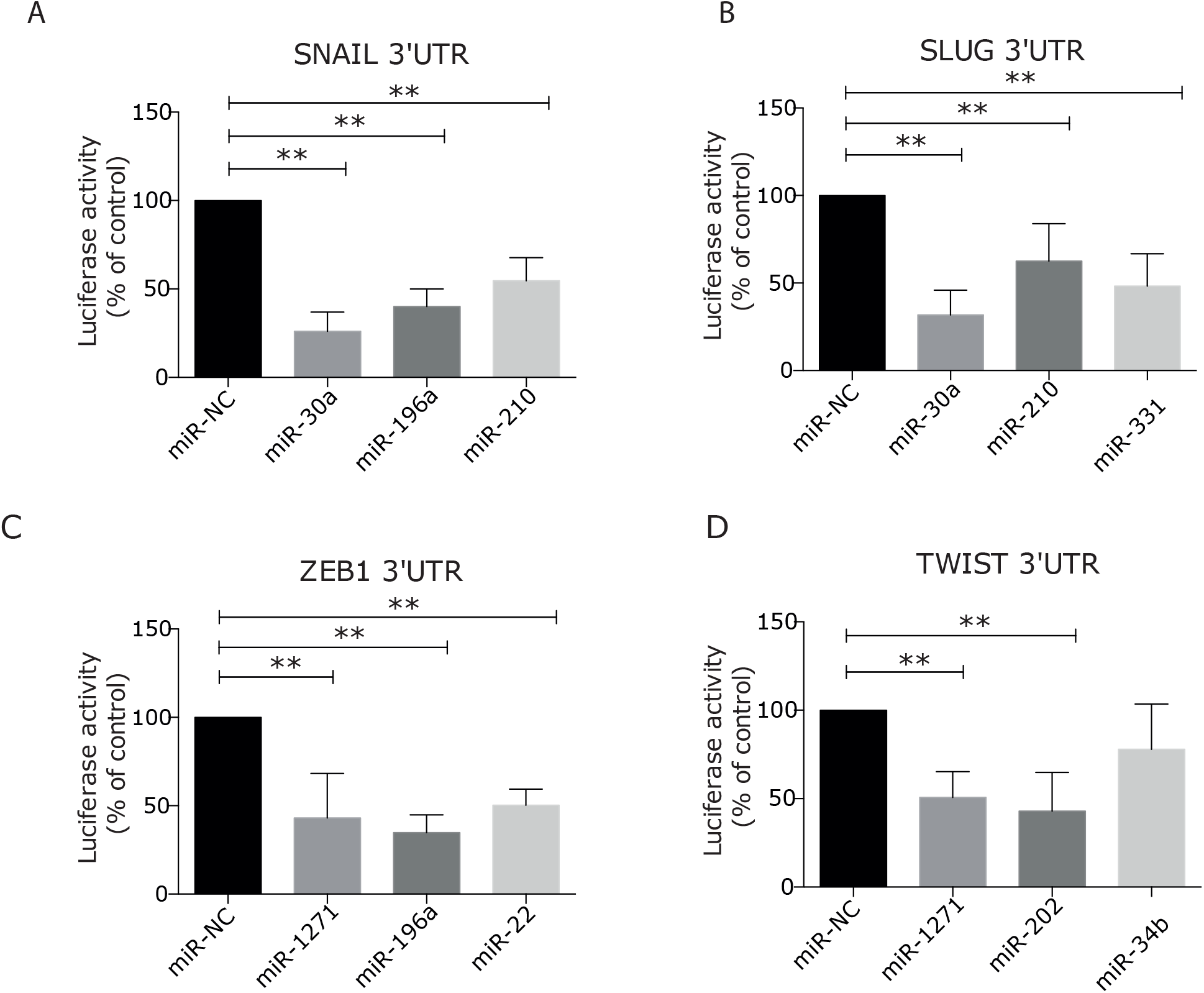
Selected microRNAs regulates SNAIL, SLUG, ZEB1 and TWIST 3’UTR. pMIR-REPORT vector with the 3’UTR of SNAIL (A), SLUG (B), ZEB1 (C) or TWIST (D) was co-transfected with the selected microRNAs at 50 nM in HEK293T cells to perform luciferase reporter assay. MiR-30a and miR-1271 were used as positive controls and a random sequence microRNA (miR-NC) was used as negative control. **, p<0.01.

### 3.3. Selected microRNAs decrease migration, invasion and cell proliferation in MDA-MB-231 cell line

During EMT, tumor cells in culture acquire the ability to migrate and invade as a previous step to metastasis [33]. We tested the capacity of the selected microRNAs, to regulate cell migration and invasion, using miR-30a and miR-1271 as controls [31,32]. Transfection of miR-210, miR-22, miR-331 and miR-34b decreased migration capacity of MDA-MB-231 cells close to 50% (Fig.2A). In addition, all evaluated microRNAs decreased cell invasion, indicating that they can modify the aptness of cancer cells to degrade a matrix to invade (Fig.2B). To our knowledge, these results demonstrate for the first time that miR-202, miR-210, miR-331 and miR-34b-3p are migration and invasion suppressor microRNAs in a breast cancer metastatic cell line.

**Fig 2.**
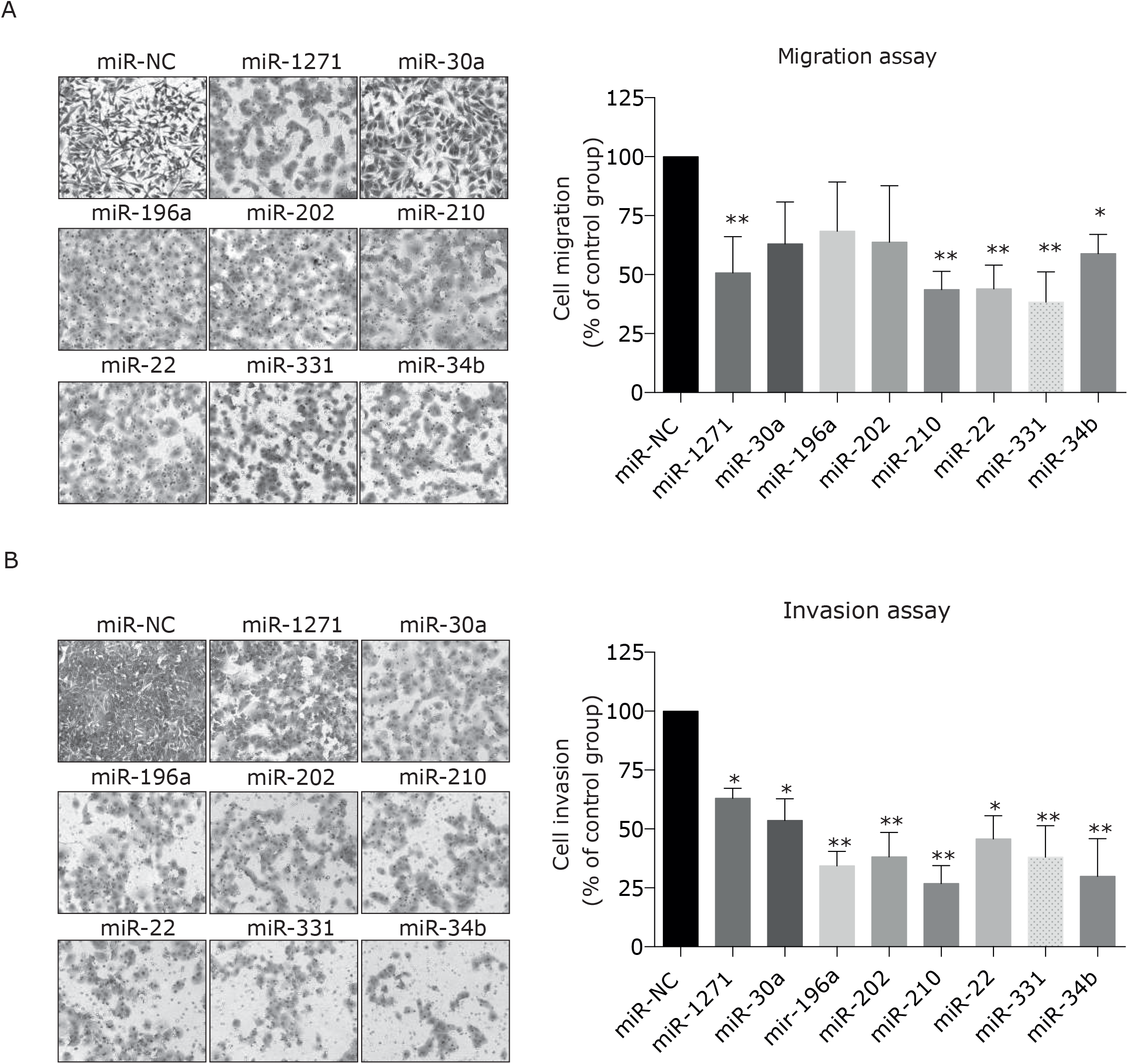
MicroRNAs decrease the metastatic traits of MDA-MB-231 cell line. Selected microRNAs were transfected in MDA-MB-231 to evaluate migration (A) and invasion (B). Cells were stained with crystal violet and five regions of each insert were analyzed using ImageJ. (C) Proliferation marker Ki67 was detected by immunofluorescence after cell transfection. (D) Colony formation capacity of transfected cells was assessed after 10 days of cell growth. Cells were stained with crystal violet and colonies formed by more than 50 cells were counted under a microscope. *p<0.05; ** p<0.01.

It is well known that microRNAs that inhibit key steps in the metastatic cascade may also repress cell proliferation [34,35]. For this reason, we evaluated the proliferation status through Ki67 marker and colony formation assays in transfected cells. MicroRNAs miR-22, miR-30a and miR-1271 decreased the proliferative potential of MDA-MB-231 cells (Fig.3A) and cell colony formation (Fig.3B) as described previously [31,36-38]. The effects of miR-196a for cell proliferation and miR-210 in colony formation have not been described previously in breast cancer, indicating that these two microRNA are novel tumor suppressor microRNAs in breast cancer. Interestingly, miR-210 only reduced the number of colonies, suggesting that this microRNA is able to control the clonogenic capacity through a mechanism distinct from proliferation.

**Fig 3.**
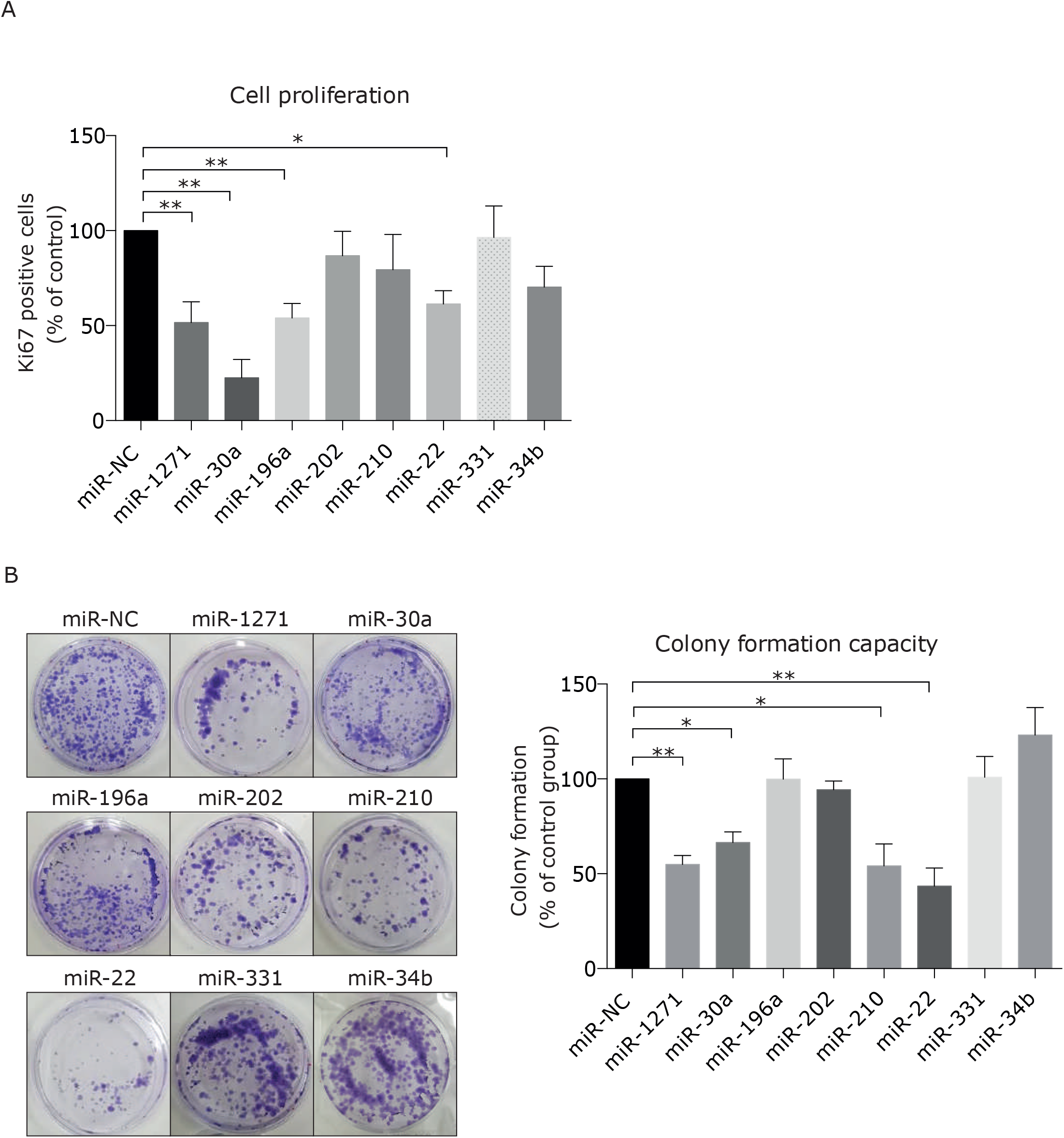
MicroRNAs decrease proliferation in MDA-MB-231 cell line. The effect of microRNAs in cell proliferation was determined through the expression of the proliferation marker Ki67 (A) and trough clonogenic assay (B). Ki67 expression was detected by immunofluorescence in transfected cells. Colony formation capacity cells was assessed after 10 days of cell growth staining the cells with crystal violet and counting colonies formed by more than 50 cells. *p<0.05; **, p<0.01.

### 3.4. miR-196a and miR-22 suppress ZEB1 expression in MDA-MB-231 cells

We transfected the microRNAs in the metastatic breast cancer cell line MDA-MB-231 to evaluate their effect on the endogenous levels of EMT transcription factors. TWIST is absent in MDA-MB-231 under the employed experimental conditions, therefore mir-202 and mir-34b were not evaluated. Neither SNAIL or SLUG levels were affected by any of the microRNAs shown in the previous experiment to inhibit luciferase activity trough their 3’UTR (Supplementary Fig.2A-B). Contrary to this, endogenous ZEB1 level decreased with transfection of miR-1271, miR-196a and miR-22 (Fig.4), revealing that these microRNAs are novel regulators of ZEB1 expression in breast cancer. Co-transfection of miR-196a and miR-22 microRNAs did not have a different effect over ZEB1 level.

**Fig 4.**
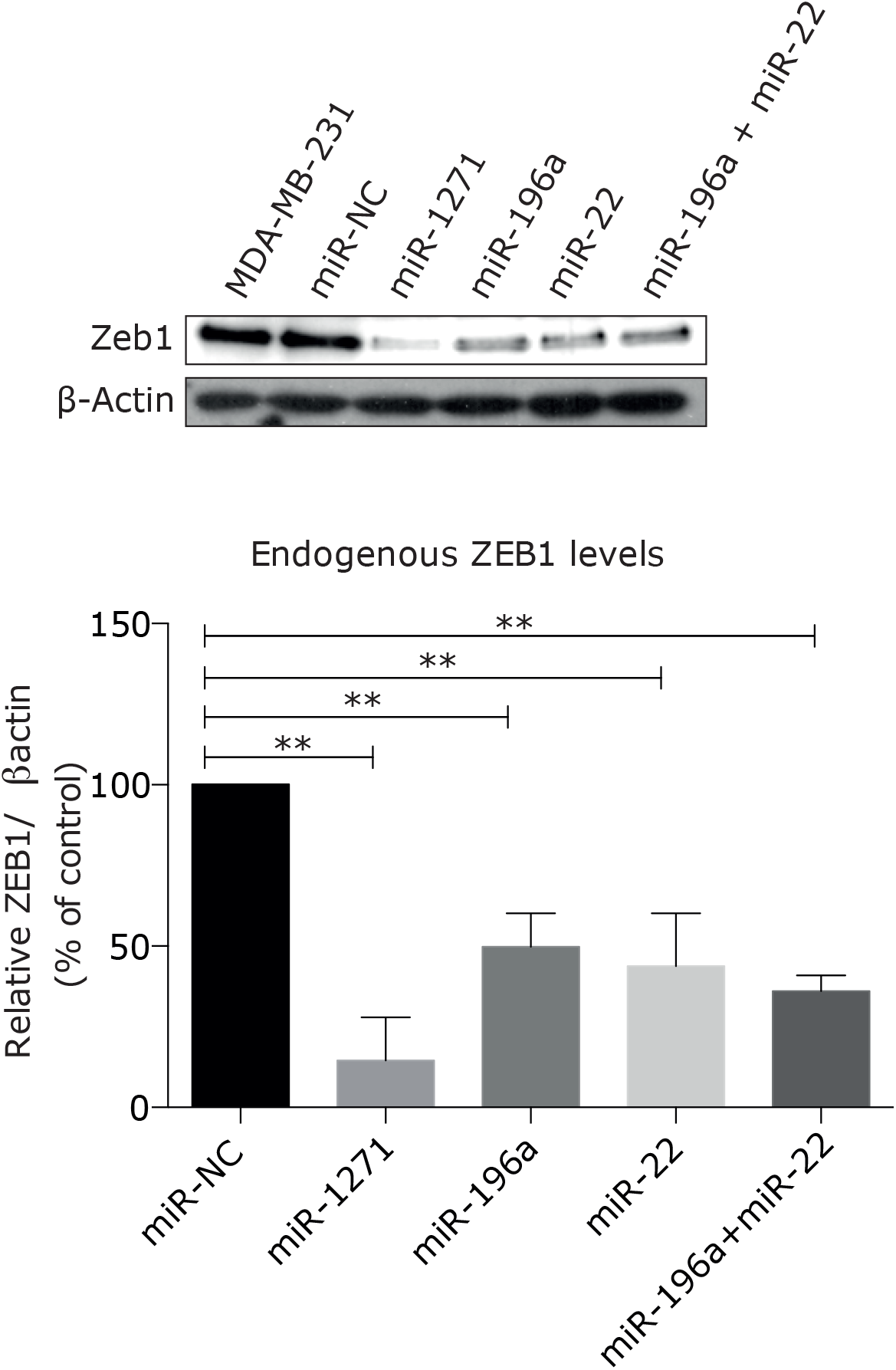
miR-1271, miR-196a and miR-22 regulates ZEB1 expression in MDA-MB-231 cell line. MiR-1271, miR-196a and miR-22 were transfected in MDA-MB-231 cells and changes in endogenous ZEB1 levels were analyzed by Western Blotting. Three independent experiments were performed and quantified using ImageJ. **, p<0.01.

### 3.5. CCR7 is another novel target of candidate microRNAs

CCR7 is a chemokine receptor which expression has been associated to lymph node metastasis in breast cancer [39,40]. Due to its relevance, we performed immunohistochemistry for CCR7 in our group of breast cancer tumors, detecting membranous stain in 96 of the 101 analyzed samples (Supplementary Fig.1B). Among tumors from patients with lymph node metastasis 42/44 were positive for CCR7 expression, but also a high proportion of non-lymph node metastasis tumors were positive for CCR7, revealing no association between the receptor expression and metastasis stage. In addition, the mean of CCR7 expression was similar between tumors with or without lymph node metastasis (H-score 148.3 vs 143.9, respectively). Interestingly, CCR7 expression was associated to tumor size (p=0.0369). These results indicate that the solely expression of CCR7 in breast cancer tissues is not a reflection of lymph node metastasis. *In silico* analysis for all studied microRNAs revealed that miR-196a, miR-202 and miR-210 were potential regulators of CCR7. We confirmed this prediction through luciferase reporter gene assays, finding that miR-196a and miR-202 decreased luciferase activity through the CCR7 3’UTR (Fig.5A). Conversely, no effect of miR-196a, miR-202 and miR-210 was detected on endogenous CCR7 level in MDA-MB-231 cells (Supplementary Fig.2C). When we tested the rest of our microRNAs, we found that transfection of miR-30a, a microRNA with no predicted *in silico* binding sites in CCR7 mRNA, decreased endogenous CCR7 expression in MDA-MB-231 cells (Fig.5B). In conclusion, we found that miR-30a is a novel regulator of CCR7 expression, and may be acting through an indirect mechanism to suppress its expression in breast cancer cells.

**Fig 5.**
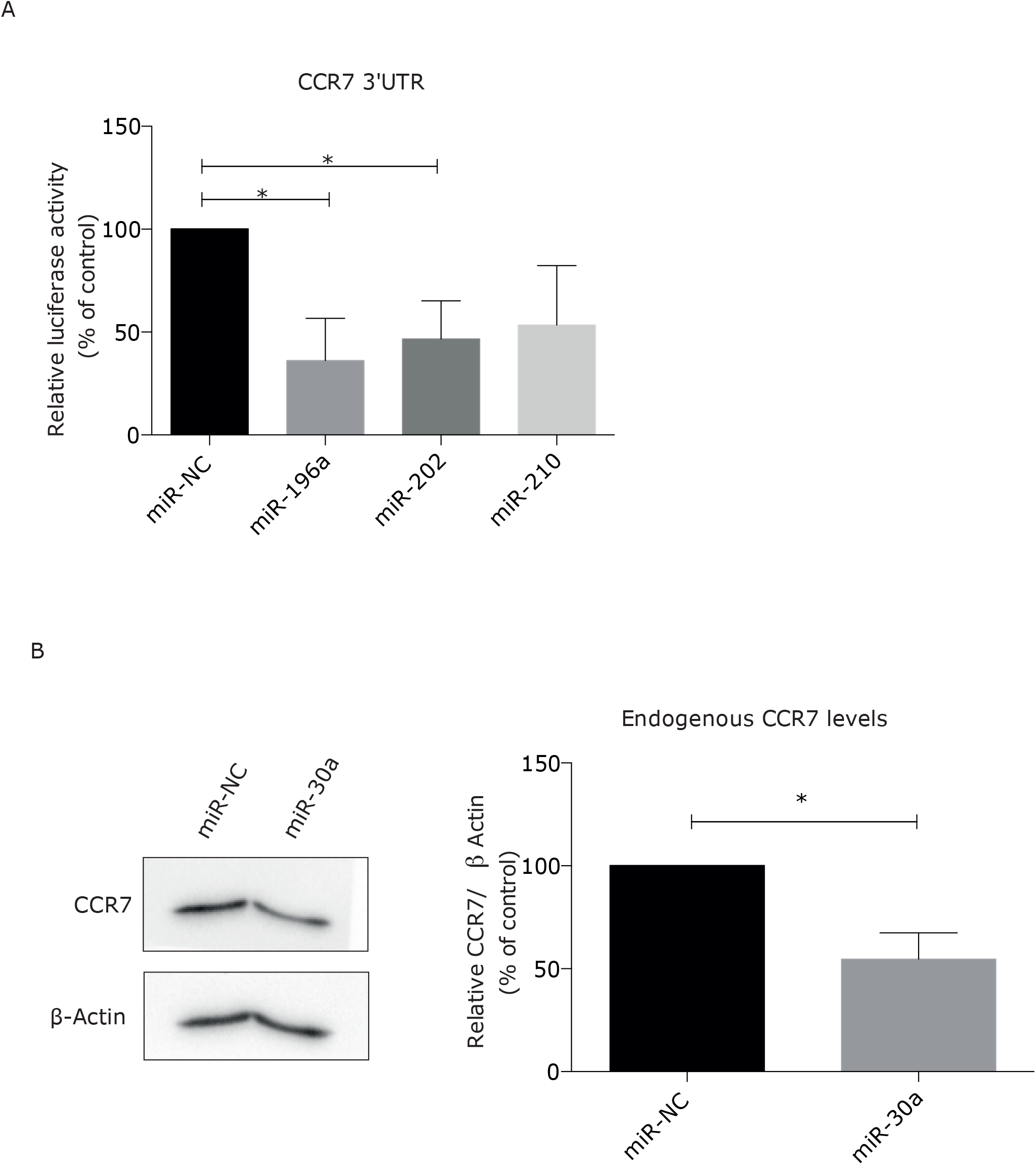
CCR7 is a novel target of regulation of miR-196a, miR-202 and miR-30a. The predicted regulation of CCR7 by miR-196a, miR-202 and miR-210 was evaluated trough luciferase reporter gene assay (A). The microRNAs were co-transfected with pMIR-REPORT luciferase vector containing the 3’UTR of CCR7 in HEK293T cells to evaluate their effect in luciferase expression. (B) CCR7 endogenous levels were assessed in MDA-MB-231 cells after transfection of miR-30a by Western Blotting. *p<0.05.

## 4. DISCUSSION

Expression of SNAIL, SLUG, ZEB1 and TWIST in primary breast tumors has been associated to lymph node metastasis [12-15]. In this study, we found expression of the four transcription factors in breast cancer tumors, but no association with lymph node metastasis was found. Expression of the EMT transcription factors is an early event in metastasis progression, inducing persistent molecular changes in tumor cells that ultimately would lead into metastasis. Because these events occur in a sequential mode, we would not expect to find a co-occurrence in a same tumor. For this reason, a lack of association between the expression of EMT transcription factors and the occurrence of metastasis, does not discard that an EMT process is occurring in breast cancer tumors. Experiments in animal models demonstrate that transient SNAIL expression is necessary to increase the development of metastasis in mice [8], suggesting that a similar scenario with temporal expression of EMT transcription factors may occur in human tumors.

MicroRNAs, which deregulation is a common feature in cancer tissues [17], are relevant modulators of metastasis through the regulation of EMT. In this study, we proved that miR-196a, miR-202, miR-210, miR-22 and miR-331 could regulate luciferase expression through their predicted 3’UTR targets. All these microRNAs are novel repressors for SNAIL, SLUG, ZEB1 and TWIST, suggesting being new potential regulators of EMT in cancer.

MiR-196a and miR-22 decreased endogenous ZEB1 levels in MDA-MB-231 cells, demonstrating for the first time that these two microRNAs are regulators of ZEB1 levels in cancer cells. The effect of both microRNAs in the luciferase reporter assays strongly suggest that this is a direct effect, acting through ZEB1 3’UTR.

In addition, mir-196a and miR-22 decreased migration, invasion and/or cell proliferation in MDA-MB-231 cells as previously described [38,41,42], suggesting that these effects may be related to the decrease of ZEB1 in MDA-MB-231 cells. MiR-22 is also a regulator of SNAIL expression [43,44], which suggest that this microRNA could be regulating EMT at least through two different targets.

Although we selected miR-1271 because of its role as ZEB1 regulator [32], this regulation was not previously described in breast cancer. In this work, we demonstrated that miR-1271 also regulates ZEB1 in breast cancer cells, reinforcing its role as a suppressor of EMT in cancer.

MicroRNAs miR-331 and miR-34b-3p, whose role in breast cancer metastasis has not been previously studied, decreased invasion and migration of MDA-MB-231 cells. Their effect on these two crucial steps in metastasis development, reinforces their role as metastasis suppressors described in lung and cervical cancer [45,46].

MiR-210 is a microRNA that has been principally studied under hypoxic conditions in cancer. Our results revealed that overexpression of miR-210 decreased invasion and migration of MDA-MB-231 cells, suggesting being a repressor of EMT. MiR-210 also decreased cell colony formation without affecting the potential of cell proliferation. We suggest that this microRNA could decrease the formation of colonies by a mechanism different from proliferation, such as apoptosis or cell detachment [47].

The relevance of CCR7 expression in the tumor cell membrane for lymph node metastasis led us to evaluate the expression of this receptor in breast cancer tumors. As for EMT transcription factors, no association was found between CCR7 expression and the presence of lymph node metastasis, suggesting that CCR7 expression by itself cannot predict the occurrence of lymph node metastasis in patients. In relation to microRNAs in lymph node metastasis, few studies have described a differential expression in tumors in relation to lymph node status [24-27]. Only let-7a has been described as a CCR7 regulator in breast cancer [48]. In this sense, this paper describes additional three microRNAs as potential regulators of CCR7 through different approaches: mir-196a, mir-202 and mir-30a. In our group of tumors, miR-30a is decreased in samples with lymph node metastasis and in tumors with high expression of CCR7. Because miR-30a can regulate invasion, cell proliferation and CCR7 expression in breast cancer cells, downregulation of miR-30a in breast cancer tumors could be relevant for the development of lymph node metastasis. Taking together our results, we suggest that loss of microRNA expression in breast cancer tumors could promote metastasis through the expression of EMT inducers as SNAIL, SLUG, ZEB1 or TWIST, and the enhancement of metastatic traits of cancer cells. These microRNAs may also control the development of lymph node metastasis through the regulation of CCR7. In this sense, reestablishing the expression of microRNAs as miR-196a, miR-22 and miR-30a could represent a strategy to treat metastatic breast cancer.

## Acknowledgements

We thanks to M.E. Andrés and V. Torres for providing the cell lines for the study, P. Gajardo and M.I. Gómez for technical support, L. Fernández and C. Llanos for donation of reagents, C. Álvarez for critically reading the manuscript. This work was supported by CONICYT # 21151349.

## Conflict of interest

The authors declare no conflict of interests.

---

**Supplementary Table 1.** Clinicopathological features of the 101 breast cancer tumors analyzed.

| **Feature** | **N** | **%** |
| --- | --- | --- |
| **Primary Tumor (T)** |  |  |
| T1 | 25 | 24.8 |
| T2 | 64 | 63.3 |
| T3 | 9 | 8.9 |
| T4 | 3 | 3.0 |
| **Regional Lymph Nodes (N)** |  |  |
| N0 | 46 | 45.5 |
| N1 | 36 | 35.6 |
| N2 | 13 | 12.9 |
| N3 | 6 | 6.0 |
| **Differentiation** |  |  |
| Grade 1 | 18 | 17.8 |
| Grade 2 | 34 | 33.7 |
| Grade 3 | 31 | 30.7 |
| **Tumor type** |  |  |
| Invasive ductal carcinoma | 82 | 81.2 |
| In situ ductal carcinoma | 2 | 2.0 |
| Invasive lobular carcinoma | 13 | 12.9 |
| Medullary | 1 | 1.0 |
| Metaplastic | 2 | 2.0 |
| No information | 1 | 1.0 |
| **Tumor subtype** |  |  |
| Luminal A | 46 | 45.5 |
| Luminal B | 15 | 14.9 |
| Luminal/HER2 | 16 | 15.8 |
| HER2 | 4 | 4.0 |
| Triple negative | 18 | 17.8 |
| Not classified | 2 | 2.0 |

**Supplementary Figure 1.**
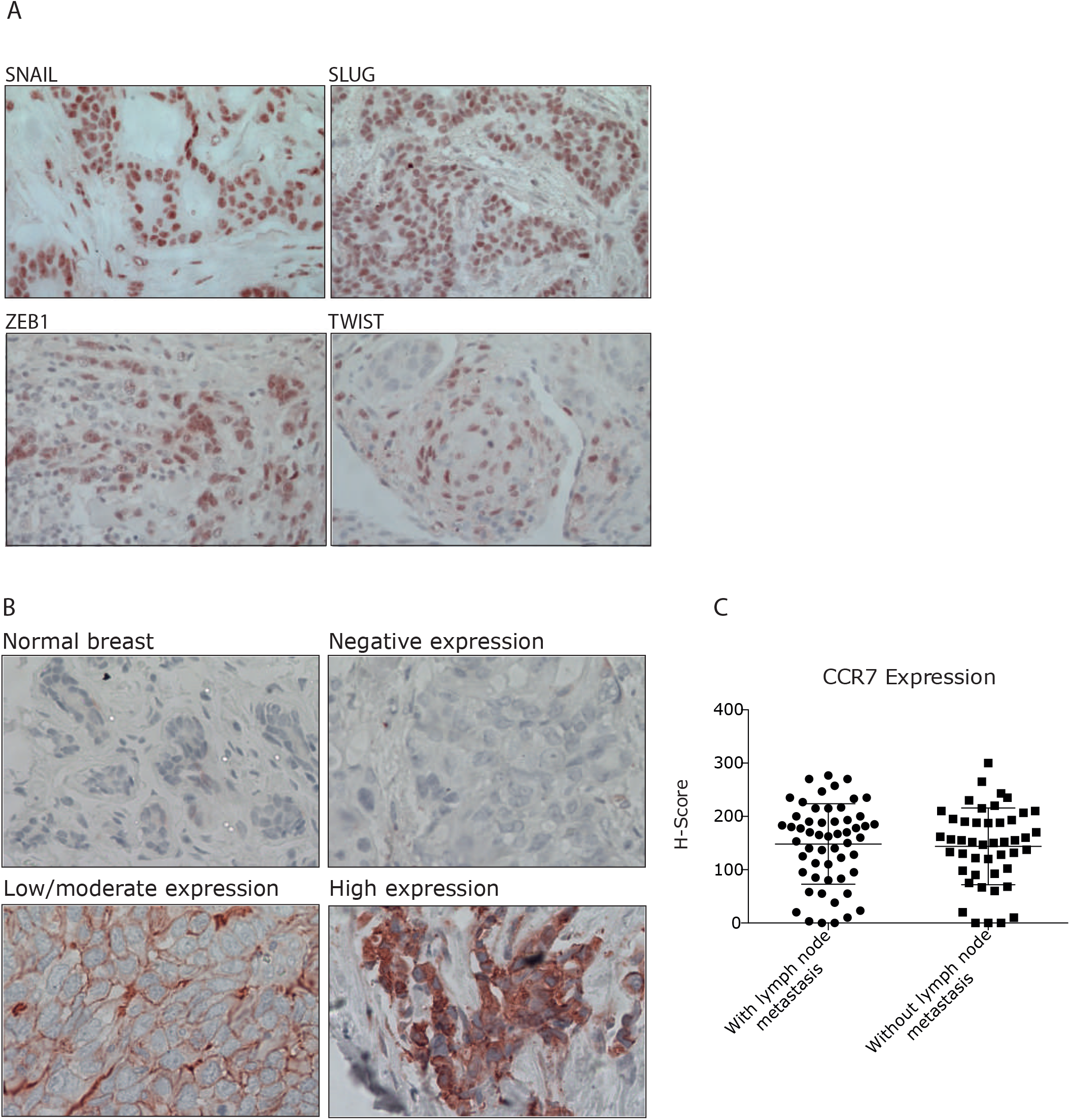
Immunohistochemistry in breast cancer tissues. (A) SNAIL, SLUG, ZEB1 and TWIST were detected in breast cancer tissues by immunohistochemistry. The four analyzed transcription factors presented nuclear stain. Images at 40X. (B) CCR7 was detected in normal breast and breast cancer tissues by immunohistochemistry. H-Score was calculated for every tumor, considering membranous stain. Mean expression (H-Score=146) was used as cut-off to define high or low CCR7 expression. Images at 40X. (C) CCR7 expression in breast cancer tumors with lymph node metastasis versus tumors without lymph node metastasis. There is a high variability of CCR7 expression in breast cancer tumors, and the media of expression does not differ between both groups.

**Supplementary Figure 2.**
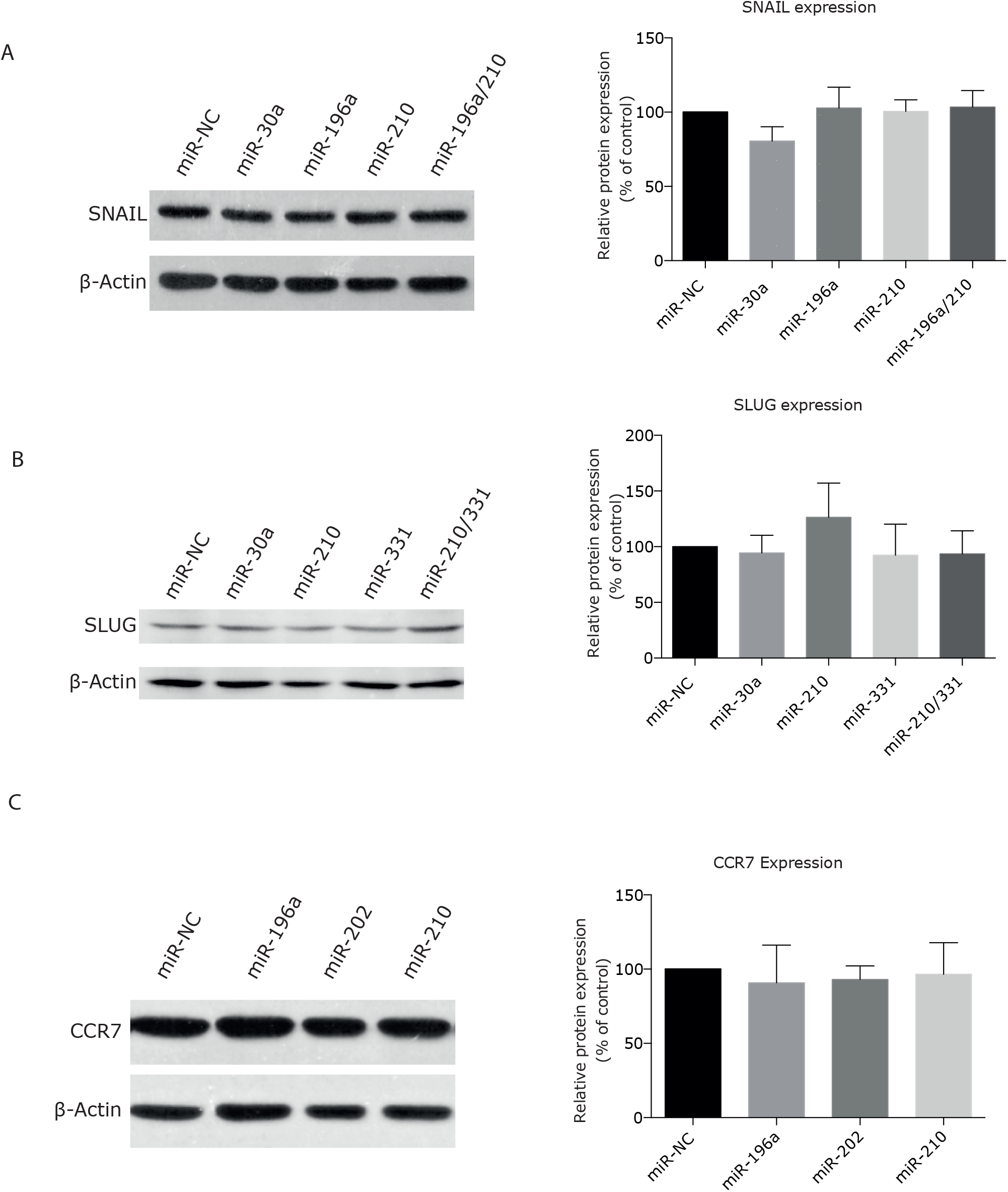
Effect of microRNAs in endogenous SNAIL, SLUG and CCR7 levels on MDA-MB-23 cell line. Cells were transfected with selected microRNAs and changes in endogenous SNAIL (A), SLUG (B) and CCR7 (C) levels were analyzed by Western Blotting. Three independent experiments were performed and quantified using ImageJ.

## Notes

### Competing Interest Statement

The authors have declared no competing interest.

